# Human immunodeficiency reveals GIMAP5 as lymphocyte-specific regulator of senescence

**DOI:** 10.1101/2021.02.22.432146

**Authors:** Ann Y. Park, Michael Leney-Greene, Matthew Lynberg, Xijin Xu, Lixin Zheng, Yu Zhang, Helen Matthews, Brittany Chao, Aaron Morawski, Ping Jiang, Jahnavi Aluri, Elif Karakoc Aydiner, Ayca Kiykim, John Pascall, Isil Barlan, Sinan Sari, Geoff Butcher, V. Koneti Rao, Richard Lifton, Safa Baris, Ahmet Ozen, Silvia Vilarinho, Helen Su, Michael J. Lenardo

## Abstract

Elucidating the molecular basis of immunodeficiency diseases is a powerful approach to discovering new immunoregulatory pathways in humans. Here we report 10 affected individuals from 4 families with a new immunodeficiency disease comprising of severe progressive lymphopenia, autoimmunity, immunodeficiency, and liver disease due to recessive loss of function variants in *“GTPase of immunity-associated proteins” protein 5 (GIMAP5)*. We show that the disease involves the progressive loss of naïve T lymphocytes and a corresponding increase in antigen-experienced, but poorly functional and replicatively senescent T cells. In vivo treatment of Gimap5-deficient mice with rapamycin (an inhibitor of mTORC1) significantly restores the fraction of naïve T lymphocytes. Furthermore, a GIMAP5-deficient human patient who was treated with rapamycin (sirolimus) showed a remarkable reduction in spleen/lymph node size. Together, these observations reveal that GIMAP5 plays a critical role in lymphocyte metabolism which is essential for senescence prevention and immune competence, suggesting that an inhibitor of mTORC1 could be a valuable clinical intervention in treating patients deficient for GIMAP5.

## Introduction

Guanosine triphosphate (GTP) hydrolases (G proteins) are a large class of regulatory switches for internal cellular functions found in all living organisms (*1, 2*). The “GTPase of immunity-associated proteins” (GIMAPs; previously “immune-associated nucleotide-binding” proteins: IANs) are proteins containing an AIG type of GTP binding domain, highly conserved among vertebrates and higher plants and apparently descended from a primordial gene, that are now found in tandem clusters of GIMAP paralogues in the metazoan genome (*3*). In vertebrates, the GIMAPs are mainly expressed in cells of the immune system (*4*). In humans, there are 7 translated GIMAP genes and one pseudogene (GIMAP3) linked on chromosome 7q36.1. Each human GIMAP protein differs in its intracellular distribution implying that each may have a unique metabolic role during immune responses (*5, 6*).

GIMAP5 is best known as one of the causative genes in the classic Biobreeding (BB) rat model of type 1 diabetes (*7*). Genetic GIMAP5 deficiency in BB rats causes markedly decreased T and natural killer (NK) cell longevity leading to type I diabetes and colitis (*7, 8*). Subsequently, several independently-derived mouse strains deficient for GIMAP5 all having a common phenotype of progressive loss of T and NK longevity affecting CD8^+^ T cells more profoundly than CD4^+^ T cells, concomitant with normal B cell counts; increased granulocytes; anemia; colitis; and liver failure with hepatocellular degeneration and increased extramedullary hematopoiesis(*9–11*). The lymphocyte disorder and liver defects appear to be cell-intrinsic (*9–11*). In humans, single nucleotide polymorphisms in GIMAP genes are linked to type 1 diabetes, systemic lupus erythematosus, and asthma (*12–14*). One lymphopenic/thrombocytopenic patient with splenomegaly was reported to have a GIMAP5 point mutation leading to protein loss but the data were limited to only a single subject and did not validate the causative nature of the GIMAP5 allele (*15*).

The metabolism of lymphocytes, especially T cells, is highly regulated to adapt to specific molecular needs during activation, proliferation, survival and exhaustion (*16, 17*). Activated effector T lymphocytes use glycolysis to provide energy and metabolites required to synthesize macromolecules for cell expansion, whereas naïve T cells rely on oxidative phosphorylation and fatty acid oxidation (*17*). The mechanistic target of rapamycin (mTOR) kinase and its associated regulatory factors comprising the mTOR Complex 1 (mTORC1) is a critical nexus between these metabolic states and controls cell growth, protein synthesis, senescence and other cellular functions (*18*). As a key metabolic regulator, mTOR is an effective point of intervention for potent pharmaceuticals such as rapamycin to constrain pathological T cell immune responses (*19*).

In this study, we identify patients suffering from a novel recessive Mendelian disease of immune dysregulation comprising progressive lymphopenia, autoimmunity, immunodeficiency and liver disease due to loss of function mutations in *GIMAP5*. GIMAP5 deficiency in patients causes severe reduction in naïve T lymphocytes with an increase of senescent effector T cells. Importantly, we show that the inhibition of mTOR provides clinical benefit in both murine and human GIMAP5 deficiency.

## Results and discussion

### Characterization of patients with immune dysregulation

We identified 10 affected individuals in 4 pedigrees with lymphopenia, splenomegaly, liver disease, thrombocytopenia, lymphadenopathy, and recurrent lung infections with bronchiectasis (Figure 1A-C and Table 1; additional clinical descriptions are in Table S1). The patients experienced fatigue related to anemia, bleeding, and compromised breathing due to respiratory infections. Three affected individuals from pedigree 1 are deceased–two from infections and liver failure and one from hypoxemic respiratory failure due to pulmonary embolus (Table 1 and Figure 1A). We performed a bone marrow biopsy on one patient (P1.5) that revealed myelodysplasia with increased atypical megakaryocytes, increased erythroid progenitors, granulocytic hypoplasia but normal fractions of lymphocyte precursors implying that peripheral lymphocyte depletion was due to a defect in mature cells.

**Figure 1.**
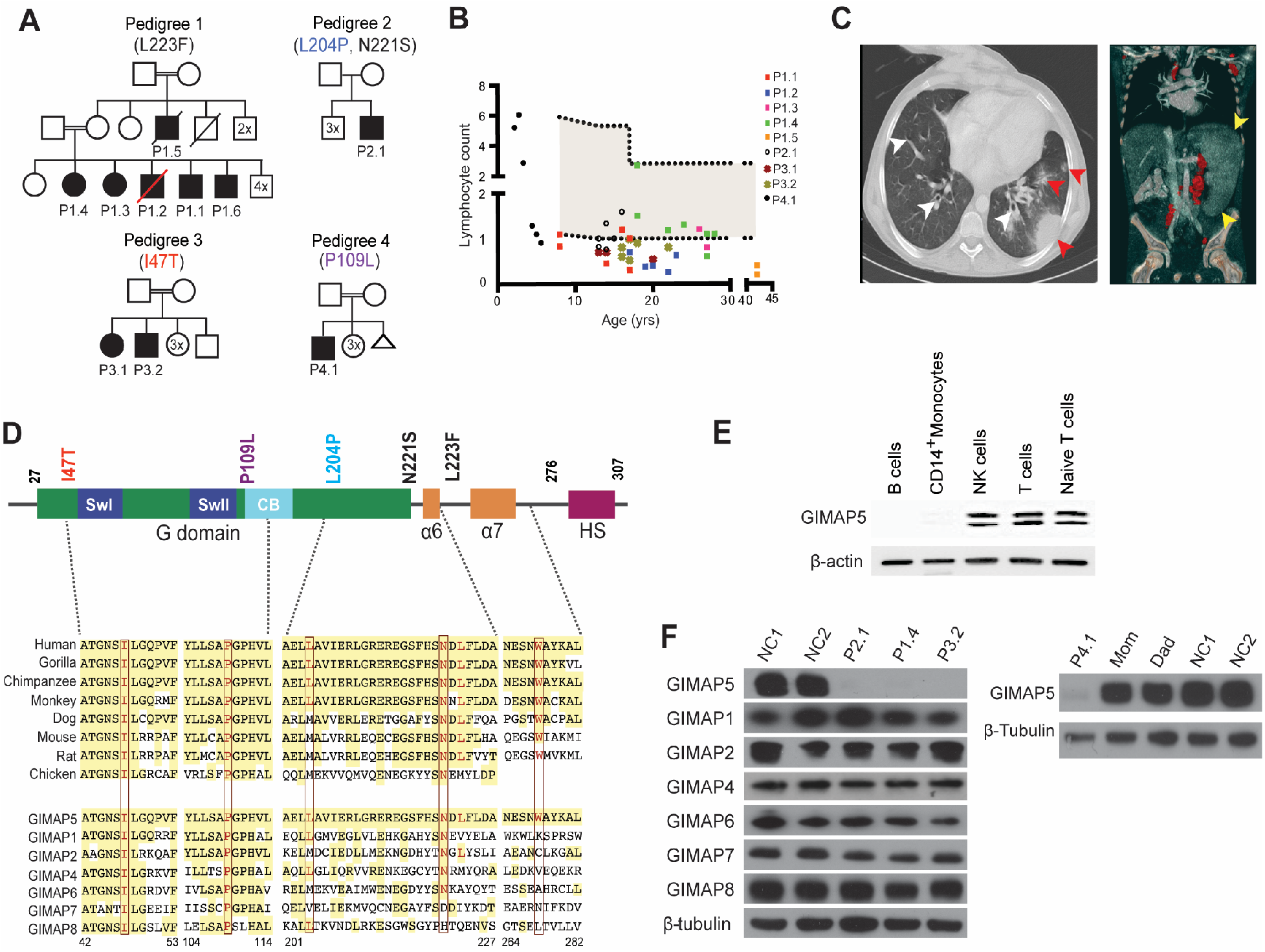
**(A)** Simplified pedigrees indicating affected individuals (filled symbols; P = patients), unaffected family members (open symbols), deceased individuals (red strikethrough); and consanguinity (double line) between parents. Additional same sex siblings are enumerated by 2x, 3x, etc. Patient amino acid changes are shown in single letter code at the respective position. Extended pedigrees are in fig. S1. **(B)** Absolute lymphocyte counts (ALC) of patients as a function of age in years (yrs). Shaded area represents the 95% confidence interval for normal individuals. **(C)**. Computerized tomography radiographs of patient P1.2 of the lungs (left) with areas of bronchiectasis with widened damaged bronchioles (white arrowheads) and opaque areas of lung consolidation due to pneumonia (red arrowheads) or the abdomen (right) illustrating enlarged spleen (yellow arrows) and enlarged lymph nodes **(D)** (Top) Schematic of the GIMAP5 protein depicting the GTPase domain, alpha helices (α) 6 and 7, and the hydrophobic segment. (Bottom) Amino acid alignment with the residue number shown at the bottom illustrating conservation in GIMAP5 orthologues (top) and 7 human paralogues (bottom). Amino acids affected by mutations are shown in red. **(E)** Immunoblot of the GIMAP5 protein in isolated primary human CD19^+^ B cells, CD14^+^ monocytes, CD56^+^ NK cells, and T cells with a β-actin loading control. Note that GIMAP5 is present as 2 isoforms in human cells. **(F)** Expression of GIMAP1, 2, 4, 5, 6, 7 and 8 proteins in patient or normal control (NC) T cells compared to a β-tubulin loading control. Note that GIMAP5 is overexposed thereby partly obscuring the two isoforms.

**Table 1.** Patient characteristics.

| Table 1. Patient Characteristics |  |  |  |  |  |  |  |  |
| --- | --- | --- | --- | --- | --- | --- | --- | --- |
| Patient |  | Immunodysregulation |  |  |  | Immunology Lab Values |  | Other |
| ID | Demographics | Splenomegaly | Lymphadenopathy | Lung Infections | Lymphopenia | Dysgammaglobulinemia | Autoantibodies | Liver Disease |
| P1.1 | Died at 17 y.o.<br>Male | Yes | Yes | Difuse bronciectasis | ↓T/B cells, Neutrophils | IgG↓ IgA↓ IgM: N | Anti thrombocyte antibodies | Yes |
| P1.2 | Died at 22 y.o.<br>Male | Yes | Yes | Focal bronchiectasis, atelectasis, ground glass opacities and infultation | ↓T/B cells, Neutrophils | IgG: N IgA: N IgM: N | Not determined | Yes |
| P1.3 | 31 y.o. Female | Yes | Not determined | Focal bronciectasis, peribroncial thickening | ↓Lymphocytes | IgG↑ IgA: N IgM: N | Not determined | Not determined |
| P1.4 | 33 y.o. Female | Yes | Yes | Consolidation at the left lung | ↓T and B cells | IgG↑ IgA↑ IgM: N | Not determined | Not determined |
| P1.5 | Died at 44 y.o.<br>Male | Yes | Yes | Focal bronchiectasis | ↓T cells, Neutrophils | IgG↑ IgA: N IgM: N | ANA(+) | Yes |
| P1.6 | 38 y.o. Male | Yes | Not determined | Not determined | ↓T cells, Neutrophils | IgG: N IgA: N IgM: N | ANA(+) | Yes |
| P2.1 | 21 y.o. Male | Yes | Yes | Ground glass changes and nodular pulmonary infiltrates | ↓Lymphocytes | IgG↓ IgA↓ IgM↑ | Anti-thrombocyte, neutrophil antibodies | Yes |
| P3.1 | 23 y.o. Female | Yes | Yes | Chronic fibrotic changes | ↓Lymphocytes | IgG↑ IgA: N IgM: N | Not determined | Yes |
| P3.2 | 25 y.o. Male | Yes | Not determined | Not determined | ↓Lymphocytes | IgG: N IgA: N IgM: N | Not determined | Yes |
| P4.1 | 7 y.o. Male | Yes | Not determined | Not determined | ↓T/B cells, Neutrophils | IgG: N IgA: N IgM↓ | Not determined | Yes |

### Identification of loss-of-function destabilizing mutations in GIMAP5

We hypothesized an autosomal recessive Mendelian disease due to the inheritance pattern and consanguinity, and exome sequencing revealed deleterious variants in all families in the GIMAP5 gene (Figure 1D and S1B and Table S2). We considered protein-altering variants with minor allele frequency <0.1% (from dbSNP, NHLBI, 1000 Genomes, Genome Aggregation Database (gnomAD) and Yale exome databases - including 894 Turkish exomes) that were homozygous or compound heterozygous and deleterious (Figure 1D and Table S2). Affected individuals from pedigree 1 were homozygous for p.L223F (c.667C>T) while P2.1 had compound heterozygous mutations causing p.L204P (c.610T>C) and p.N221S (c.662A>G) alleles. Affected individuals in pedigree 3 had a homozygous I47T (c.140T>C) substitution and the proband in pedigree 4 harbored homozygous p.P109L (c.326C>G). Unaffected siblings and parents tested were either heterozygous or wild type. We found that the altered GIMAP5 amino acids were highly conserved and the CADD and polyphen scores predict that the changes are damaging (Figure 1D and Table S2). Furthermore, the gene mutations underlying the p.I47T and p.L223F alterations have not been reported in the gnomAD database, p.P109L has been detected once, while the nucleotide changes underlying p.L204P and p.N221S have minor allele frequencies of 0.002 and 0.000012, respectively (Table S2). Parametric linkage analysis using the WES data from the parents was also performed using affected individuals and two unaffected siblings in pedigree 1. Pedigree 3 yielded a logarithm of the odds (LOD) score of 1.8, pedigree 4 yielded a LOD score of 1.2 and pedigree 1 yielded a LOD score of at least 6.2, providing a highly significant combined LOD score of 9.2 (data not shown). Finally, the phenotypes found in our cohort resembles mouse strains with *GIMAP5* loss of function mutations as well as an isolated individual with a *GIMAP5* variant and lymphopenia (*9, 10, 15*).

To understand why the disease primarily affected T and NK cells, we checked a publicly available database of microarray data from 78 different human tissues and found that GIMAP5 mRNA is predominantly expressed in T cells and NK cells (Figure S1C) (*20*). CD34^+^ progenitors or CD19^+^ B cells had no expression, which differs from mice (*5, 21*). Protein immunoblots confirmed robust expression in purified human T cells or CD56^+^ NK cells, but none in CD14^+^ monocytes and CD19^+^ B cells (Figure 1E). In our patients, we observed no or little residual GIMAP5 protein expression suggesting that the p.I47T, p.L204P, p.N221S, p.L223F and p.P109L missense amino acid substitutions are destabilizing (Figure 1F). We also found that patient T cells had normal levels of the GIMAP1, 2, 4, 6, 7 and 8 proteins indicating an independent loss of human GIMAP5. Hence, the selective defects in normal T and NK cell homeostasis are a consequence of the cell-specific expression and loss of human GIMAP5.

### T lymphocyte phenotyping of GIMAP5 deficient patients

Since the patients showed decreased T cell numbers and GIMAP5 is highly expressed in T cells, we carried out a more detailed investigation of the circulating T cell subpopulations to understand the severe immunodeficiency. We observed a marked decrease in CD45RA^+^CCR7^+^ naïve phenotype CD4^+^ and CD8^+^ T cells with a corresponding increase in CD45RA^−^CCR7^−^ effector memory T cells with little change in the CD45RA^+^CCR7^−^ central memory T cells (Figure 2A). Because the depletion of naïve T cells suggested a state of immune overactivation, we analyzed the level of CD57, a marker of immune cell post-activation senescence. Among CD8^+^ memory T cells, we observed markedly expanded CD57^+^ T cells in the patients signifying excessive proliferation and replicative senescence (Figure 2B). Replicative senescence was confirmed by shortened telomere lengths in T cells from two representative patients that were below the 1st percentile for their age (Figure 2C). These results indicate unregulated immune overactivation especially given the young age of the patients. Despite these results, we saw no abnormalities in early molecular events after T cell receptor (TCR) stimulation (Figure S2A and B). The depletion of naïve T cells and remaining senescent T cells accounts for the profound immunodeficiency of the patients.

**Figure 2.**
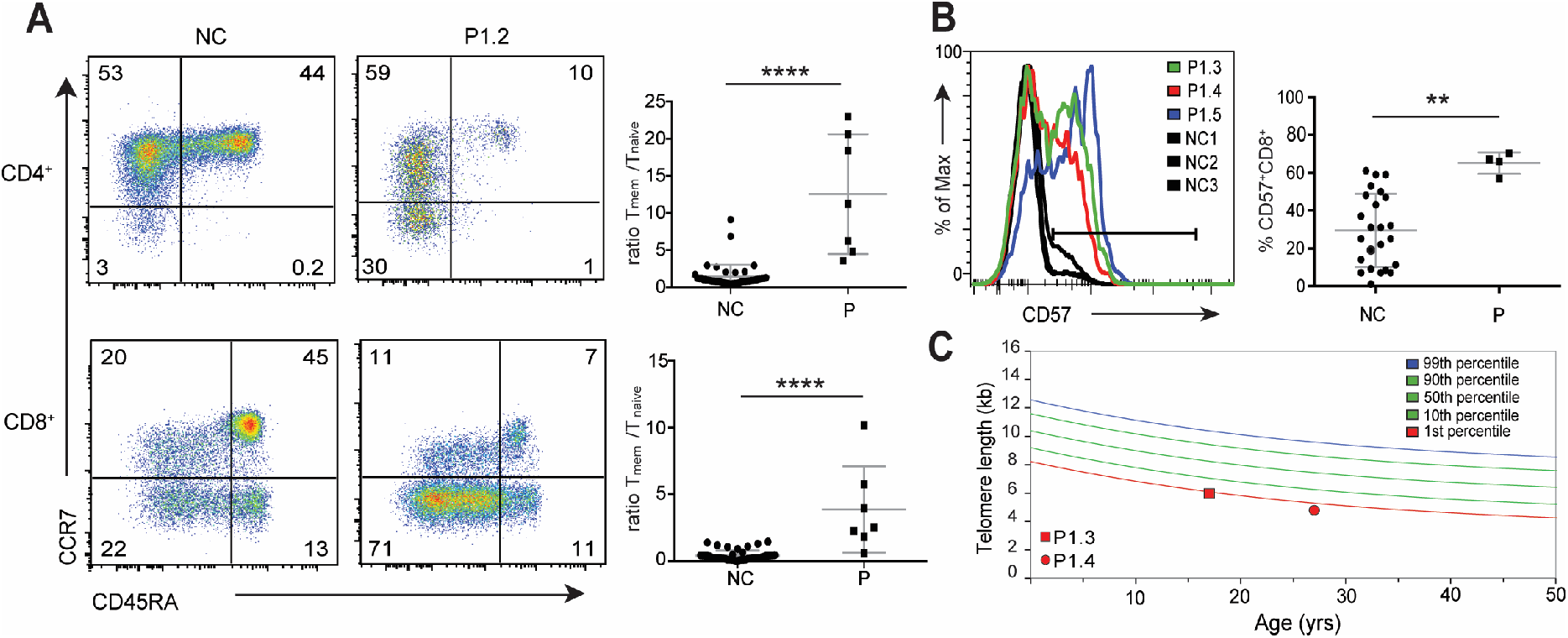
**(A)** Flow cytometry dot plots comparing a patient (P1.2) and healthy control (NC) PBMCs gated on CD3^+^ (all), CD4^+^ (top) or CD8^+^ (bottom) and stained for CD45RA and CCR7 (left). Ratio of effector/memory CD45RA^−^CCR7^−^ T cells (T_mem_) to naïve CD45RA^+^CCR7^+^ T cells (T_naïve_). **(B)** Flow cytometry histogram showing CD57 expression on patient cells (P, colors) compared to normal controls (NC, black) (left). Quantification of CD57^+^CD8^+^ T cells as defined by the indicated gate in (left). **(C)** Flow cytometric fluorescence *in situ* hybridization analysis of telomere length in kilobases (kb) within the lymphocyte population of patients P1.3 (square) and P1.4 (circle). Colored lines represent the indicated percentile isobars for telomere length versus age in years (yrs). Individual points plotted in (A) and (B) represent the mean of individual patients and pooled normal controls averaged across 1-5 independent measurements. Statistical significance was calculated by Student’s t-test **p<0.01, ****p<0.0001.

### Inhibition of mTORC1 provides clinical benefit in both murine and human GIMAP5 deficiency

Differentiation to effector phenotypes has been shown to be driven by mTORC1 during the immune response (*16*). Notably, mice carrying alleles leading to hyperactive mTORC1, including TSC1 deficient mice also suffer from peripheral lymphopenia with normal thymopoiesis and a loss of peripheral naïve T cells as Gimap5 deficient mice (GIMAP5^sph/sph^) (*22*). Similarly, human patients with gain-of function mutations in the PI3K subunit p110δ that promote hyperactivation of mTOR in T cells have a clinical phenotype that significantly overlaps with the GIMAP5-deficient patients (*19*). Furthermore, previous work has shown that T cells isolated from GIMAP5^sph/sph^ mice had increased mTORC1 activity at baseline as measured by the phosphorylation levels of pS6 at serines 235 and 236 (*23*). Therefore, we investigated if the inhibition of mTOR activity can provide beneficial effects in suppressing senescence phenotype in GIMAP5-deficient mice and patients.

We confirmed previous findings regarding increased levels of pS6 as well as increased cell size in peripheral T cells from GIMAP5^sph/sph^ mice via flow (Figure 3A). GIMAP5^sph/sph^ mice or littermate controls were injected daily with rapamycin intraperitoneally. We found that *in vivo* treatment of Gimap5^sph/sph^ mice with rapamycin could significantly reduce both pS6 and cell size (forward scatter) to levels comparable to wild type controls (Figure 3A). Importantly, we observed that *in vivo* treatment with rapamycin significantly increased the fraction of naïve T cells present in both the spleen and the blood (Figure 3B, data not shown). Furthermore, P3.1 was treated with sirolimus (an inhibitor of mTORC1) beginning in November of 2013 and continuing up to the present and we observed a remarkable reduction in spleen/lymph node size (Figure 3C). In this same time period his severe psoriasis was similarly resolved. Taken together, these data suggest that mTORC1 inhibitors may be a valuable clinical intervention in treating human patients deficient for GIMAP5.

**Figure 3.**
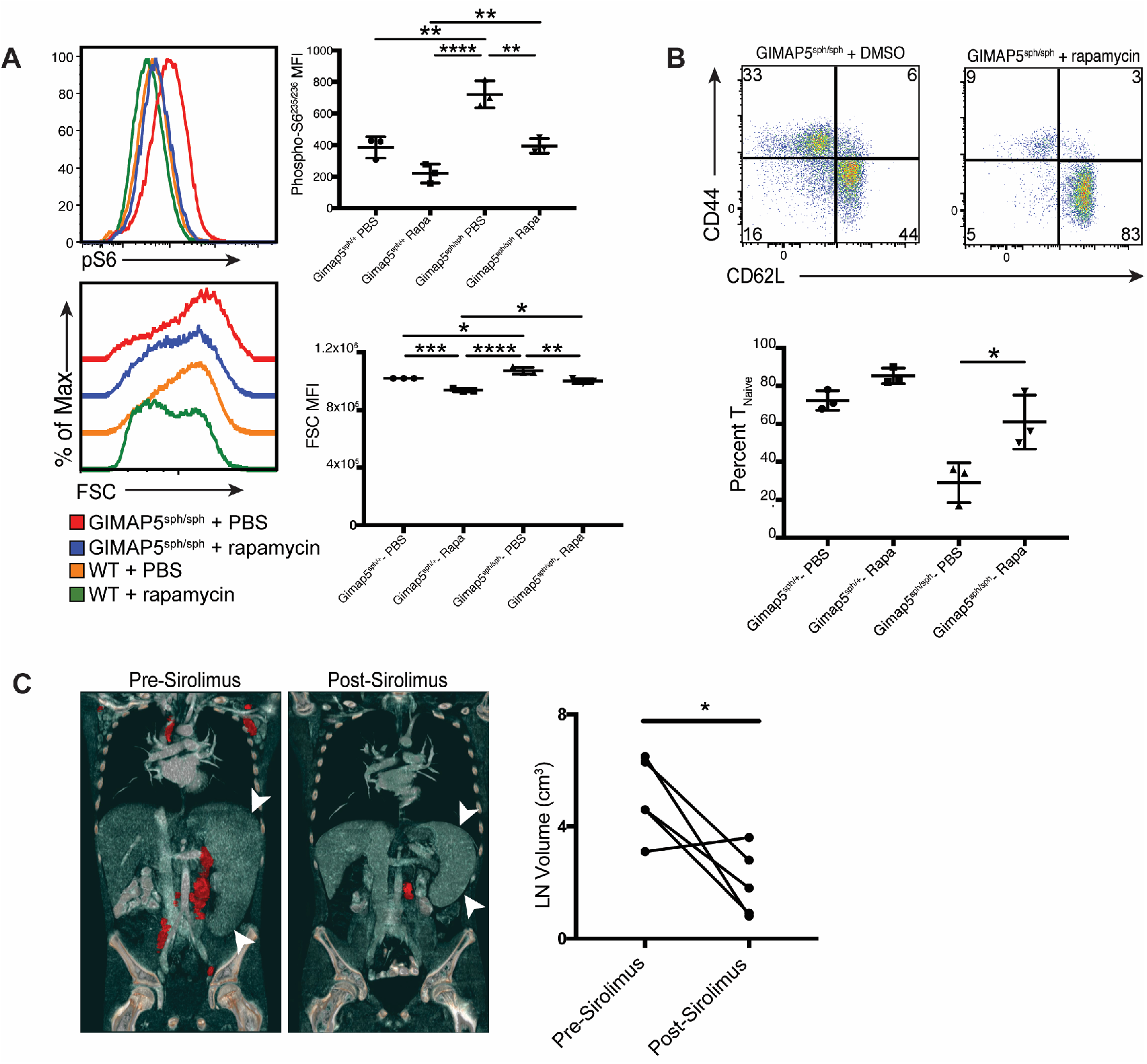
**(A)** GIMAP5^sph/sph^ mice or littermate controls were treated with rapamycin as described in the methods section and then readouts of mTORC1 activity (size, pS6^235/236^) were measured in T cells via flow cytometry. **(B)** Mice were treated with rapamycin as in (A) and then the fraction of naïve cells in the spleen was measured. **(C)** Abdominal CT scans of a GIMAP5-deficient patient pre/post sirolimus treatment for 2 years. Three dimensional reconstruction was used to measure the volume of 5 lymph nodes. Data shown in A-B is representative of three experiments, with significance calculated by ANOVA *p<0.05, **p<0.01, ***p<0.001, ****p<0.0001. Data in C is representative of 3 separate experiments, while statistical significance was done via Student’s t test in D.

In conclusion, we have identified a novel human immunodeficiency we now term “GISELL Disease” for **<u>G</u>**IMAP5 defects with **<u>I</u>**nfections, **<u>S</u>**plenomegaly, **<u>E</u>**nlarged lymph nodes, **<u>L</u>**ymphopenia and **<u>L</u>**iver nodular regenerative disease. GISELL Disease is caused by loss of function mutations in *GIMAP5* that involves progressive loss of naïve T lymphocytes and an increase in senescent T cells. Importantly, *in vivo* treatment with rapamycin significantly restored the fraction of naïve T lymphocytes and improved clinical course. Furthermore, these observations show that GIMAP5 is essential for human immunity by preventing lymphocyte senescence, suggesting a role of GIMAP5 in the regulation of lymphocyte metabolism.

## Materials and methods

### Human subjects

All human subjects (or their legal guardians) in this study provided written informed consent in accordance with Helsinki principles for enrollment in research protocols that were approved by the Institutional Review Board of the National Institute of Allergy and Infectious Diseases, National Institutes of Health (NIH). Blood from healthy donors was obtained at the NIH Clinical Center under approved protocols. Mutations will be automatically archived by Online Mendelian inheritance in Man (OMIM) once the paper is published.

### Mice

All protocols were approved by the NIAID Animal Care and Use Committee and followed NIH guidelines for using animals in intramural research. Gimap5sph/sph mice were a kind gift from Dr. Kasper Hoebe (Janssen Research and Development Spring House, Spring House, Pennsylvania, USA). Littermate C57BL/6 mice were used as wild-type controls. Gender-matched 6- to 12-week-old mice were used for all experiments described in this study.

### Primary T cell culture

Human PBMCs were isolated by Ficoll-Paque PLUS (GE Healthcare) density gradient centrifugation, washed twice in PBS, and resuspended at 106/mL in complete RPMI 1640 (cRPMI) medium containing 10% FBS, 2 mM glutamine, and 100 U/mL each of penicillin and streptomycin (Invitrogen). Cells were activated with 1 μg/mL anti-CD3 (clone HIT3α, BioLegend) and 1 μg/mL anti-CD28 (clone CD28.2, BioLegend). After 3 days, activated T cells were washed and then cultured in cRPMI with 100 U/mL Recombinant Human IL-2 (rhIL-2, R&D). In the case of patient cells, freshly isolated or frozen PBMCs were stimulated at 5×106/mL in cRPMI with 2 μg/mL of PHA-L (lectin from red kidney bean (Phaseolus vulgaris), Sigma-Aldrich L2769), in the presence of 1 μg/mL anti-CD28 and 1 ng/mL recombinant IL7 (BioLegend) for 48 hours. The cells were washed twice with cRPMI and resuspended in media at 1 × 106/mL with 100 IU/mL Recombinant Human IL2 and cultured for up to 3 weeks with fresh rhIL2 and medium supplemented every 2 days.

### Exome and Whole Genome Sequencing Analysis

Genomic DNA was isolated from PBMCs of proband, parents and healthy relatives from each pedigree. Exome sequencing were generated using SureSelect Human All Exon 50Mb Kit (Agilent Technologies) coupled with Illumina HiSeq sequencing system (Illumina). Whole Genome Sequencing (WGS) were generated based on Standard Coverage Human WGS from Broad Institutes. For individual samples, WES produced ~50-100X sequence coverage for targeted regions and WGS produced 60X coverage for proband and 30X coverage for family members. DNA sequence data was aligned to the reference human genome (build 19) using Burrows-Wheeler Aligner (BWA) with default parameters and variants were called using the Genome Analysis ToolKit (GATK) (Li and Durbin, 2009). Variants were then annotated by functional impacts on encoded proteins and prioritized based on potential disease-causing genetic model. Variants with minor allele frequency < 0.1% in the dbSNP (version 137), 1000 Genomes (1,094 subjects of various ethnicities; May 2011 data release), Exome Sequencing Project (ESP, 4,300 European and 2,203 African-American subjects; last accessed August 2016), ExAC databases (61,000 subjects of various ethnicities; March 2016 data release) or Yale internal database (2,500 European subjects) were filtered. Autosomal-recessive inheritance was investigated and genes with rare homozygous or compound heterozygous variants were prioritized.

### Flow Cytometry

For standard surface staining, PBMCs and T cell blasts were washed once in FACS buffer and stained in 50 μl of FACS buffer containing indicated fluorochorome-labeled antibodies for 30 mins on ice. Cells were then washed three times in FACS buffer and fixed before acquisition on either a LSR II or LSRFortessa (BD Biosciences). Data was analyzed using FlowJo v. 10.5.3.

### Telomere analysis

Frozen PBMCs from patient P1.3 and P1.4 were shipped to Repeat Diagnostics for measurement of telomere length via Flow-FISH. A fluorescently labeled nucleic acid probe was hybridized with the TTAGGG repeats in telomeres. Signal strength was then plotted relative to telomere lengths for healthy control subjects in order to calculate the percentile for each patient.

### Orthologues and Other Human GIMAPs

Full-length orthologues of GIMAP5 protein sequences from several species and related human GIMAP-family members (GIMAP1, 2, 4, 6, 7 and 8) were obtained from GenBank. Protein sequences were aligned using the Clustal Omega algorithm.

### Immunoblotting

Protein lysates were isolated from cell pellets via a 15 minute incubation in lysis buffer (1% NP-40, 150mM sodium chloride, 50mM Tris-HCl, pH 7.4 plus Halt™ Protease and Phosphatase Inhibitor Cocktail) and then clarified via centrifugation at 14,000 to 16,000 × g at 4°C for 15 minutes. Total protein levels were measured by BCA assay (Pierce) prior to reduction with 5% beta-mercaptoethanol and SDS Loading Solution (Quality Biological) at 95°C for 10 minutes. Equal amounts of protein were electrophoresed on either 4-12% Bis-Tris or 10% Bis-Tris gels (NuPAGE, Life Technologies) and wet-transferred to nitrocellulose membranes. These were then blocked in 5% non-fat milk in TBST (0.05% Tween-20) and probed with primary antibodies to GIMAP1, GIMAP4, GIMAP5, GIMAP6 and GIMAP7 (generous gifts from Dr. Geoffrey Butcher), GIMAP2 and GIMAP8 (produced by GenScript), β-Actin (Abcam). Binding of primary antibodies was then measured via HRP-tagged antibodies specific for the relevant species (SouthernBiotech) and visualized by chemiluminescent substrate (Immobilon Classico/ Forte western HRP substrate or Thermo Fisher SuperSignal West Dura Extended Duration Substrate).

### Calcium flux assessment

T cell blasts from healthy donors and patients were loaded with Indo-1-AM dye (Invitrogen) at a final concentration of 0.5 μM with PowerLoad (Thermo Fisher) in HBSS with 20 mM HEPES for 20 mins at room temperature. Flux was assessed by flow cytometry upon restimulation with anti-CD3 (HIT3α, BioLegend) plus protein A (Sigma).

### *In Vivo* Rapamycin treatments

Rapamycin (LC labs) was dissolved in DMSO to generate a concentrated stock solution. This was then further diluted in PBS for in vivo use. 7-9 week old age and sex matched mice were injected intraperitoneally with 2mg/kg of rapamycin daily for two weeks prior to isolation of tissues for further experimentation.

## Acknowledgments

The authors would like to thank the referring physicians as well as our patients and families who participated in this study; the University of Washington Center for Mendelian Genomics and all contributors to Geno2MP for use of data included in Geno2MP.

## Funding section

This work was supported by the Division of Intramural Research, National Institute of Allergy and Infectious Diseases, NIH; the German Research Foundation (SFB958, project A12).

