## Supplemental Data for "Human immunodeficiency reveals GIMAP5 as lymphocyte-specific regulator of senescence"

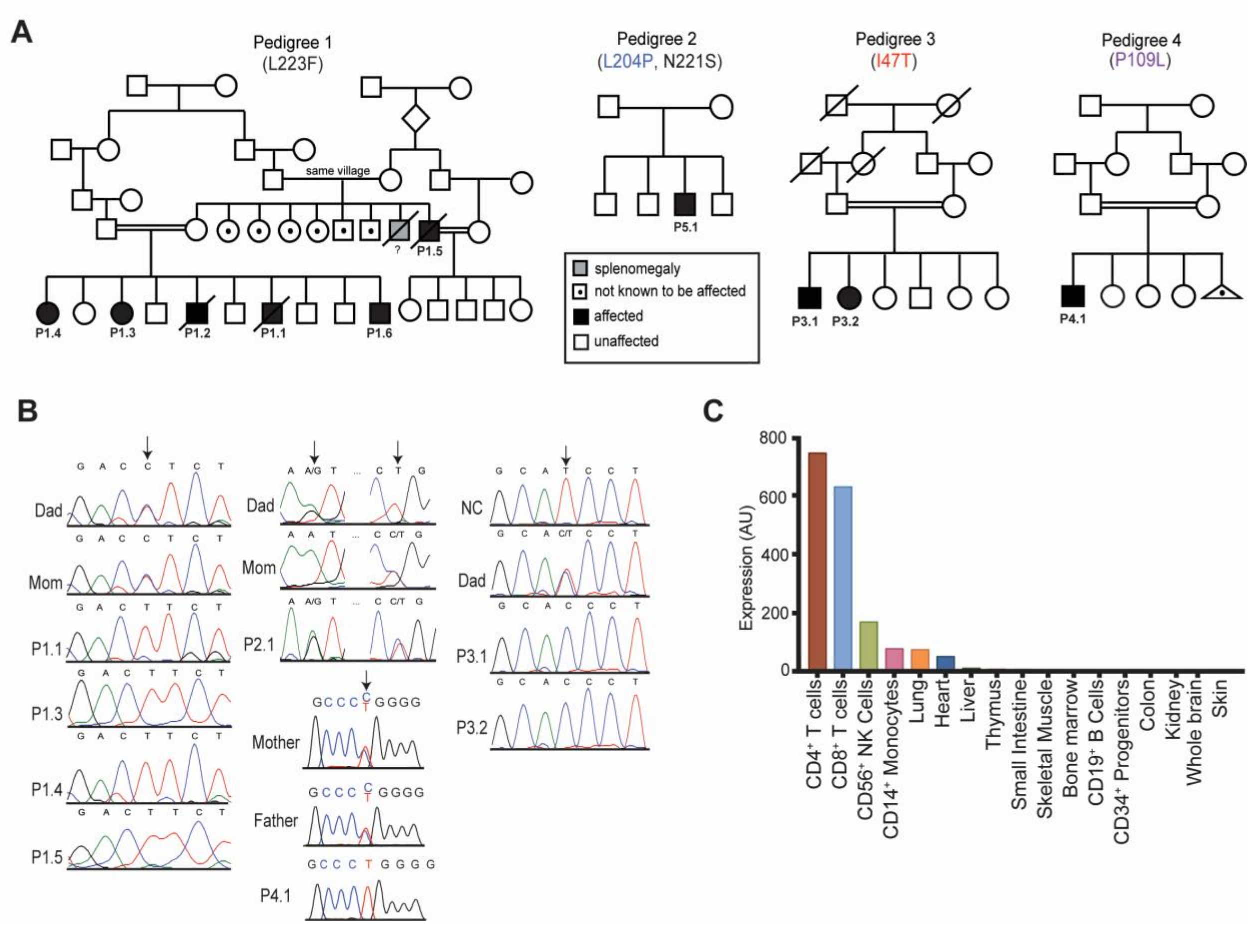


**Figure S1. In depth patient pedigrees and characterization of GIMAP5 mutations**

**(A)** Extended pedigree information for each kindred.

**(B)** Sanger sequencing confirmation of GIMAP5 mutations from affected individuals as well as healthy family members.

**(C)** Microarray RNA expression of GIMAP5 displayed in descending order of abundance in human tissues. Shown are the 22 highest expressing tissues of 79 tested. Modified from (Su et al., 2004).

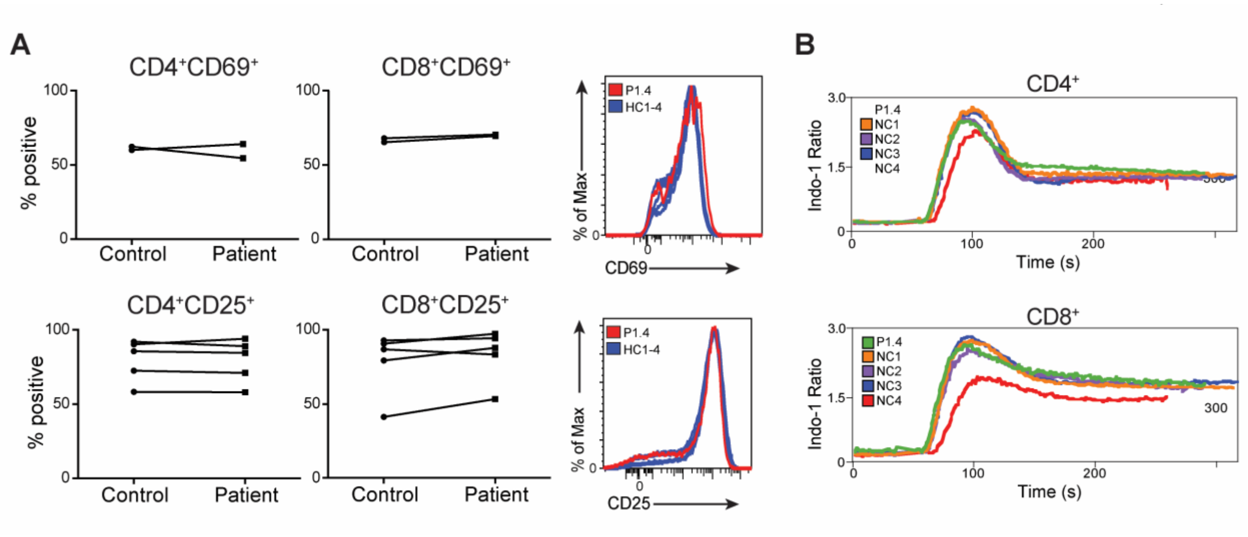


**Figure S2. Activation of primary T lymphocytes is normal in our patient cohort**
(A) Levels of T cell activation markers (CD25 and CD69) on T lymphocytes in PBMCs isolated from healthy controls or patients 48 hours after activation.
(B) Measurements of calcium flux levels following acute restimulation of cycling human T cell blasts from healthy controls or patients.

### Supplementary Table 1. Summary of the clinical features of patients with bi-allelic *GIMAP5* mutations

| | Clinical and Laboratory Findings | | | | | | | | | | |
|---|---|---|---|---|---|---|---|---|---|---|---|
| | P1.1 | P1.2 | P1.3 | P1.4 | P1.5 | P1.6 | P2.1 | P3.1 | P3.2 | P4.1 | |
| **Immunologic Alterations** | Thrombocytopenia, Neutropenia, Anemia, Lymphopenia | Thrombocytopenia, Neutropenia, Anemia, Lymphopenia | Thrombocytopenia, Anemia, Lymphopenia | Anemia, Lymphopenia | Thrombocytopenia, Neutropenia, Anemia, Lymphopenia | Thrombocytopenia, Lymphopenia | Thrombocytopenia, Neutropenia | Thrombocytopenia, Anemia, Lymphopenia | Thrombocytopenia, Lymphopenia | Thrombocytopenia, Neutropenia, Anemia, Lymphopenia | |
| **Laboratory Evaluation** | **P1.1** | **P1.2** | **P1.3** | **P1.4** | **P1.5** | **P1.6** | **P5.1** | **P3.1** | **P3.2** | **P4.1** | **Reference** |
| Lymphocyte (/µl) | **830** | **380** | 1100 | **800** | **300** | **900** | 1330 | **540** | **800** | **850** | 950-3070 |
| Neutrophil (/µl) | **300** | **110** | 2100 | 3100 | **700** | 4400 | **1000** | 2500 | 2200 | 1850 | 1560-6450 |
| Hb (g/dl) | **9** | **8.7** | **10** | **11.1** | **7.5** | 14.9 | 13.1 | 11.7 | 13.6 | **11.5** | 11.6-16.6 |
| Thrombocyte (/µl) | 350000 | **690000** | 250000 | 1640000 | **9000** | **19000** | 115000 | **123000** | **74000** | **127000** | 150000-450000 |
| CD3+ (%) | 67 | 62 | 69 | 76 | 75 | 75 | 81 | - | - | 68 | 56-84 |
| CD3+CD4+ (%) | **23** | **13** | 39 | 37 | 48 | 32 | 48 | - | - | 45 | 30-60 |
| CD3+CD8+ (%) | 44 | 48 | 28 | 30 | 28 | 43 | 30 | - | - | 18 | 11-37 |
| CD19+ (%) | **22** | **20** | 19 | 9 | 7 | 14 | 9 | - | - | **26** | 6-19 |
| CD3-CD16+/CD56+ (%) | 7 | 6 | 5 | 7 | 5 | 7 | 5 | - | - | 6 | 3-20 |
| IgA (mg/dl) | **39** | 176 | 246 | **596** | 260 | **450** | **18** | 310 | 190 | **63.5** | 139-378 |
| IgG (mg/dl) | **360** | 1340 | 1610 | **3360** | 1720 | **2060** | **380*** | 1700 | 1520 | 928 | 913-1884 |
| IgM (mg/dl) | 118 | 178 | 215 | 156 | 246 | 140 | **874** | 90 | **59** | **10.5** | 88-322 |
| | **P1.1** | **P1.2** | **P1.3** | **P1.4** | **P1.5** | **P1.6** | **P5.1** | **P3.1** | **P3.2** | **P4.1** | |
| **Infections** | Bronchiectasis, Pneumonia, Skin infections, Lymphadenitis, Candida, Acinetobacter septicemia | Bronchiectasis, Gingivitis, Thrush | None | Bronchiectasis | Bronchiectasis | None | Liver pulmonary infiltrates | None | None | Respiratory tract infection | |
| **Other Clinical Features** | Liver disease, Pulmonary embolus | Liver disease, Recurrent diarrhea Atelctasis Ground glass opacities | None | Recurrent cough, Nasal drainage | Liver disease | Liver disease | Liver disease | Liver disease | Liver disease | Liver disease | |

* post rituxan ** measured while on IVIG
**Blue**=Lower than reference **Red**=Higher than reference

**Supplementary Table 2.** Characterization of GIMAP5 mutations

| Kindred # | Chr:position (hg19) | Amino acid change | Polychen-2 (Prediction) | CADD | NHLBI | 1000G | Turkish DB | gnomAD (overall) | gnaonAD (highest frequency) |
|---|---|---|---|---|---|---|---|---|---|
| 1 | 7:150439894 | L233F[#] | 0.999 (D) | 22.7 | 0 | 0 | 0 | 0 | 0 |
| 2 | 7:150439838 | L204P[D] | 0.946 (D) | 23.0 | 0.003605 | 0.001 | 0.00168 | 0.002130 | 0.003024 (European, Non-Finish) |
| 2 | 7:150439889 | N221S[D] | 0.996 (D) | 16.35 | 0 | 0 | 0 | 1.220e-0.5 | 6.498e-05 (South Asian) |
| 3 | 7:150439367 | I47T[#] | 1.0 (D) | 25.9 | 0 | 0 | 0 | 0 | 0 |
| 4 | 7:150439553 | P109L[#] | 1.0 (D) | 27.7 | 0 | 0 | 0 | 4.067e-06 | 8.979e-06 (European, Non-Finish) |
